# Evidence for spatiotemporal shift in demersal fisheries management priority areas in the western Mediterranean

**DOI:** 10.1101/2021.02.02.429345

**Authors:** Iosu Paradinas, Joan Giménez, David Conesa, Antonio López-Quílez, Maria Grazia Pennino

## Abstract

Marine Protected Areas (MPAs) are a promising management tool for the conservation and recovery of marine ecosystems, as well as fisheries management. MPAs are generally established as permanent closures but marine systems are dynamic, which has generated debate in favour of more dynamic designs. As a consequence, the identification of priority areas should assess their persistence in space and time. Here, we develop a step-by-step approach to assess the spatiotemporal dynamics of fisheries management priority areas using standard fishery-independent survey data. To do so, we fit Bayesian hierarchical spatiotemporal SDM models to different commercially important demersal species and use the resulting maps to fit different spatial prioritisation configurations. We use these results to assess the spatiotemporal dynamics of fisheries priority areas. The proposed method is illustrated through a western Mediterranean case study using fishery-independent trawl survey data on six commercially important species collected over 17 years. We identified two fisheries priority area patterns in the study area, each predominant during a different time-period of the study, asserting the importance of regularly re-assessing MPA designs.

## 1 Introduction

The near future expects a large-scale implementation of Marine protected areas (MPAs). Different international agreements concur on protecting 10% of coastal and marine areas (CBD Aichi target 11, UN sustainable development goal 14, EU common fisheries policy target 14.5, etc.) and nearly 200 governments committed to meet this goal by 2020 (Tittensor et al. 2014). Post-2020 global conservation targets have increased this target to 30% by 2030 and to 50% by 2050 (O’Leary et al. 2016, Dinerstein et al. 2019, Baillie & Zhang 2018).

MPAs are a promising management tool for the conservation and recovery of marine ecosystems (Leenhardt et al. 2015, Giakoumi et al. 2017), as well as for fisheries management (Kaiser et al. 2007, Di Franco et al. 2016, Petza et al. 2017, Fraschetti et al. 2018, Petza et al. 2021). MPAs protect fish stocks in the no-take zone, promote greater reproductive output (Kaiser et al. 2007) and therefore sustain density dependent spill-over (Di Lorenzo et al. 2016) enhancing fisheries catches in the buffer zone and outside the MPAs (Hilborn et al. 2004, Kerwath et al. 2013, Ovando et al. 2016)

MPAs are generally established as permanent closures (Game et al. 2009), but marine systems are dynamic in space and time (Halpern et al. 2015, Gordon et al. 2018, Kroeker et al. 2020), which has generated arguments in favor of more dynamic and adaptive MPA designs (Grafton & Kompas 2005, Hughes et al. 2007, Hobday & Hartmann 2006). Therefore, the assessment of new MPAS should quantify the spatiotemporal dynamics of priority areas.

A common way of identifying multiple species priority areas is to fit spatial prioritisation algorithms using species distribution maps. The quality and resolution of species distribution maps vary from the most basic species range maps, based on presence data and/or expert opinion, to the more sophisticated species distribution maps produced by species distribution models (SDMs). SDMs are particularly useful to conservation biology (Rodríguez et al. 2007) and the selection of protected areas (Seo et al. 2009). Fishery-independent surveys collect high quality data using carefully designed protocols. Most fishery-independent survey data share common features: georeferenced data; performed periodically in the same area; collection of abundance data per species and length class; and species-specific datasets commonly have large proportions of zeros. Despite these common features, methods applied to fishery-independent survey data range from the simplest linear model to generalized spatiotemporal additive mixed models (GAMMs) and spatiotemporal geostatistical models.

A long-standing approach in designing MPAs based on species distribution maps is to use numerical spatial prioritisation algorithms such as `Marxan` (Ball & Possingham 2000), `Zonation` (Moilanen et al. 2009), or `prioritizr` (Hanson et al. 2019), which identify priority areas that cost-effectively optimise ecological objectives based on several species-specific distribution maps and a set of user-defined conditions. In the spatiotemporal framework, users may optimise a single area solution using the full time series together or produce a set of optimised areas disaggregated by time, i.e. monthly, seasonally, yearly, etc. These results provide different information that users can employ to investigate the spatiotemporal dynamics of priority areas.

Within this context, the purpose of this study was to identify priority areas for the management of the most economically important demersal species in the western Mediterranean Geographical subarea (GSA) 06, with particular emphasis on nursery areas. The western Mediterranean fishing fleet is characterised by small vessels, multiple landing sites, and multispecies catches that sell for relatively high prices (Lizaso et al. 2020). Approximately 90% of assessed stocks are exploited above the maximum sustainable yield (MSY) limits (Raicevich et al. 2018) and it has the highest percentage of unsustainable fished stocks among the 16 major seas in the world (FAO 2000). The socioeconomic models by Sola et al. (2020) suggest that in order to achieve a MSY level of vulnerable stocks requires 80% reduction of fishing effort following the current spatial planning, which in practice seems to be unrealistic (Maynou 2014, Martín et al. 2019).

In this article we identify and assess the spatiotemporal dynamics of fisheries management priority areas based on standard fishery-independent survey data. To do so, we first apply Bayesian hierarchical spatiotemporal SDMs to different commercially important demersal species (Section 2.1). Then, we use SDM results to fit different spatial prioritisation configurations (Section 2.3) and assess the spatiotemporal dynamics of fisheries management priority areas through a number of steps.

## 2 Material and methods

The data for this study come from the EU-funded Mediterranean trawl survey (MEDITS) project (Bertrand et al. 2002) carried out from April to June between 2000 and 2016. The MEDITS uses a stratified sampling design, where strata are defined by bathymetry. Sampling stations were initially placed randomly within each stratum at the beginning of the project and trawl hauls were performed in similar geographical locations every year. This study concerns the trawlable grounds of GSA06, which borders the northern Iberian Mediterranean coast, from Cap de Creus in the north to Cabo de Palos in the South (Figure 1). The data for this study comprise six species: red mullet (*Mullus barbatus*); striped red mullet (*Mullus surmuletus*); and the shortfin squid (*Illex coindetii*); European hake (*Merluccius merluccius*); monkfish (*Lophius piscatorius*) and blackbellied monkfish (*Lophius budegassa*). European hake and both monkfish species abundances were seggregated into recruits and non-rescruits. Unfortunately, the MEDITS survey only sample the non-recruit stages of red mullet, striped red mullet, and squid. Table 1 summarises the data comprised in the study.

**Table 1:** Overall presence probability and conditional-to-presence catch quantiles per species and life stage in the western Mediterranean MEDITS survey data from 2000 to 2016.

| Species | Life stage | Presence probability | $Q_{0.25}$ | $Q_{0.5}$ | $Q_{0.75}$ |
| --- | --- | --- | --- | --- | --- |
| *Mullus barbatus* | Non-recruits | 0.55 | 5 | 18 | 45 |
| *Mullus surmuletus* | Non-recruits | 0.38 | 1 | 2 | 6 |
| *Illex coindetii* | Non-recruits | 0.66 | 3 | 9 | 30 |
| *Merluccius merluccius* | Recruits | 0.73 | 14 | 47 | 106 |
| *Merluccius merluccius* | Non-recruits | 0.77 | 2 | 5 | 10 |
| *Lophius budegassa* | Recruits | 0.20 | 1 | 1 | 2 |
| *Lophius budegassa* | Non-recruits | 0.55 | 1 | 2 | 5 |
| *Lophius piscatorius* | Recruits | 0.12 | 1 | 1 | 2 |
| *Lophius piscatorius* | Non-recruits | 0.15 | 1 | 1 | 1 |

**Figure 1:**
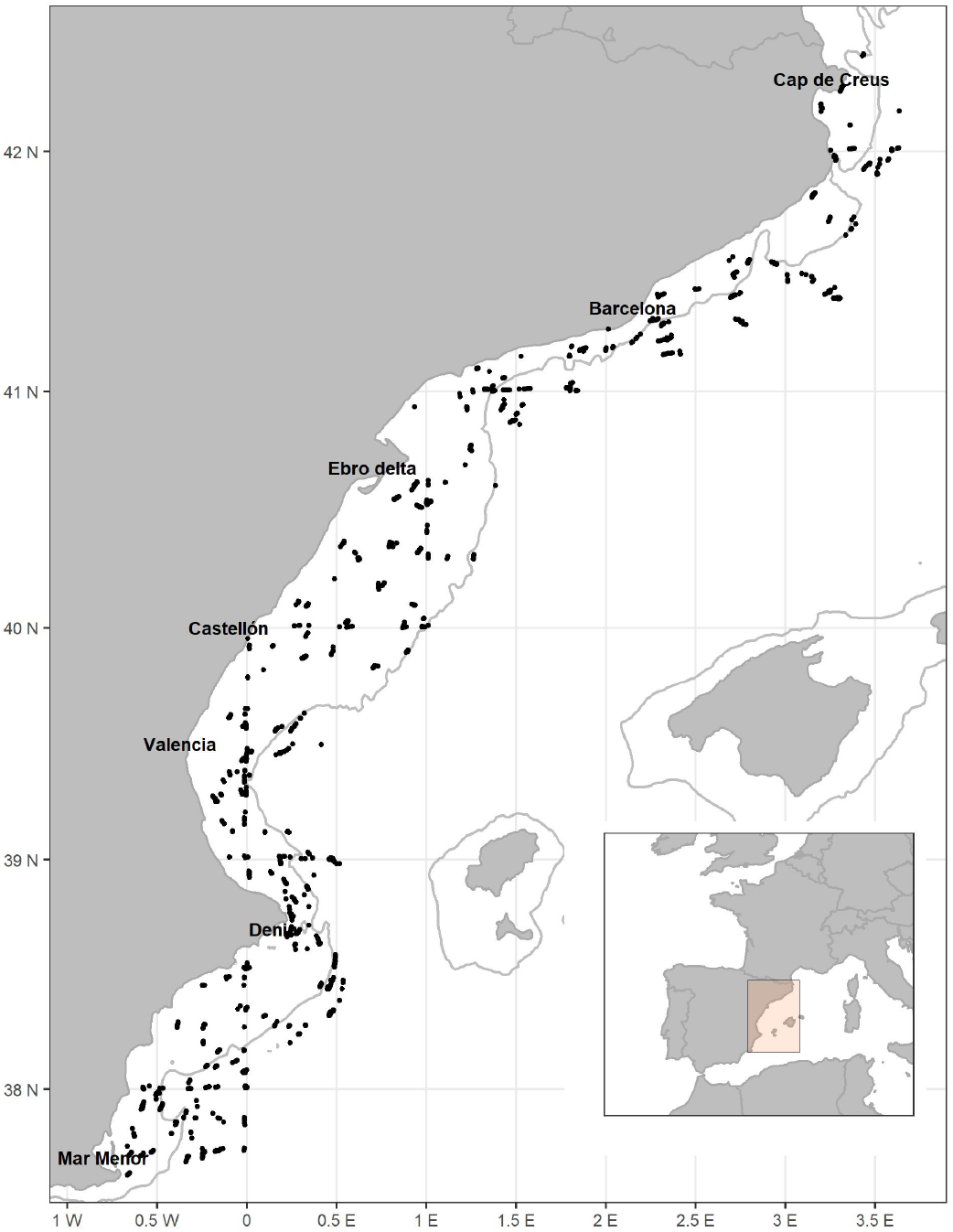
Sampling locations (black dots) of the MEDITS surveys (2000-2016). Bathy-metric lines indicate the 200 metre isobaths.

As above mentioned, our method combines spatiotemporal SDM models (Section 2.1) and spatial prioritisation algorithms (Section 2.3) to produce a set of informative results (Section 3) to assist policy makers on the design of MPAs using standard fishery-independent survey data.

### 2.1 Spatiotemporal fisheries SDMs

In order to deal with zero catch observations and the interaction between space and time, we fitted Bayesian spatiotemporal two-part or hurdle models. Hurdle models break the data in two and fit separate models to the occurrence and the abundances (Stefánsson 1996, Maunder & Punt 2004, Martin et al. 2005). Our proposed spatiotemporal structure comprised a geostatistical spatial field that evolved through a first order autoregressive temporal effect. Autoregressive effects contain a correlation parameter, namely *ρ*, that infer level of correlation or similarity between subsequent spatial distributions. Therefore, *ρ* provides important information on the degree of persistence in the process. The closer the *ρ* value is to one, the more temporally persistent the process (i.e., very high correlation between subsequent years), whereas *ρ* values closer to zero suggest more opportunistic distributions (i.e., uncorrelated distributions). See (Paradinas et al. 2017, Martínez-Minaya et al. 2018, Paradinas et al. 2020, Izquierdo et al. 2021) for further information on persistent, progressive, and opportunistic spatiotemporal fish distributions.

Our spatiotemporal proposal also included a marginal temporal trend and a non-linear bathymetric explanatory effect, both fitted through second order Random Walks (RW2), which perform as Bayesian smoothing splines (Fahrmeir & Lang 2001, Lang & Brezger 2004). RW2 models do not allow shape constrains, and therefore we visually confirmed that fitted bathymetric effects were sensible. In particular, if *Z*_*s,t*_ and *Y*_*s,t*_ are respectively the occurrence and the abundance at location *s* and time *t*, our proposed model can be formulated as:

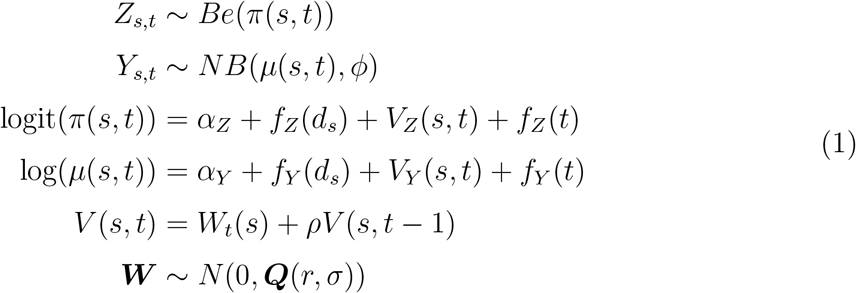

where *Be* and *NB* stand for the Bernoulli and Negative binomial distributions, respectively; *π*(*s, t*) represents the probability of occurrence at location *s* at time *t*; and *µ*(*s, t*) and *ϕ* are the mean and variance of the conditional-to-presence abundance. The linear predictors, which contain the effects linked with the parameters *π*(*s, t*) and *µ*(*s, t*) include: *α*_*Z*_ and *α*_*Y*_, terms that represent the intercepts of each respective variable; *f* (), RW2 functions with hyperparameter *γ* representing the variance of the RW2 model parametrised as unknown values *f* = (*f*_0_, …, *f*_*i−*1_)^*T*^ at *i* = 25 equidistant values of *d*_*s*_ for the bathymetry and at *i* = 17 year values of *t* for the marginal temporal trend. *V* (*s, t*) refers to a spatiotemporal field that evolves through time given the correlation parameter *ρ*. Finally, ***W*** is a geostatistical field with a covariance function defined by range *r* and standard deviation *σ*. Note that the *Be* and *NB* processes present different *V* (*s, t*) fields, *V*_*Z*_(*s, t*) and *V*_*Z*_(*s, t*), respectively.

We used the INLA package for R (Martins et al. 2013) that allows relatively fast spatial and spatiotemporal modeling (Lindgren et al. 2011). The Bayesian approach requires specification of the prior distributions for the parameters and hyperparameters of the model. We used R-INLA default vague prior distributions for the dispersion of the conditional-to-presence abundance and the fixed effects. The hyperparameters of the spatiotemporal fields and the *γ* hyperparameters of the second order random walks were set using PC priors as described by (Simpson et al. 2017) and (Fuglstad et al. 2019). Specifically, we used PC priors that followed the following criteria: a) the probability that the spatial effect range was smaller than 150 km was 0.15, to avoid very small spatial autocorrelation ranges, b) the probability that the spatial effect variance was greater than 1 was 0.20, to avoid masking the bathymetric effect through the spatial effect, and c) the probability that *γ* was greater than 0.5 in the occurrence model and greater than the observed conditional-to-presence abundance standard deviation in the conditional-to-presence model were 0.01.

### 2.2 Quantity of interest

Hurdle models provide two estimates for every location *s* and time *t*, a probability of occurrence (*π*(*s, t*)), and a conditional-to-presence abundance (*µ*(*s, t*)). While both estimates provide important information for the spatiotemporal characterisation of a species, it is common to work with the mean of a hurdle model, which can be obtained by multiplying the probability of presence and the conditional abundance (Maunder & Punt 2004, Stefánsson 1996, Zuur et al. 2009, Lecomte et al. 2013). The analytical estimation of the variance is slightly more complicated, and it varies depending on the likelihood of *µ*(*s, t*) (e.g. see (Lecomte et al. 2013) for the case of a delta-gamma and Poisson-gamma models). However, within the Bayesian paradigm, we can easily approximate hurdle model posterior distributions by resampling from the marginal distributions of *π*(*s, t*) and *µ*(*s, t*). To do so, we first predict *n*_*π*_ presence-absence values from *π*(*s, t*) and then we sample *n*_*µ*_ times from *µ*(*s, t*) (*n*_*µ*_ being the predicted number of presences), to produce our results. Figure 2 shows an example using *Mullus barbatus* results for the year 2000.

**Figure 2:**
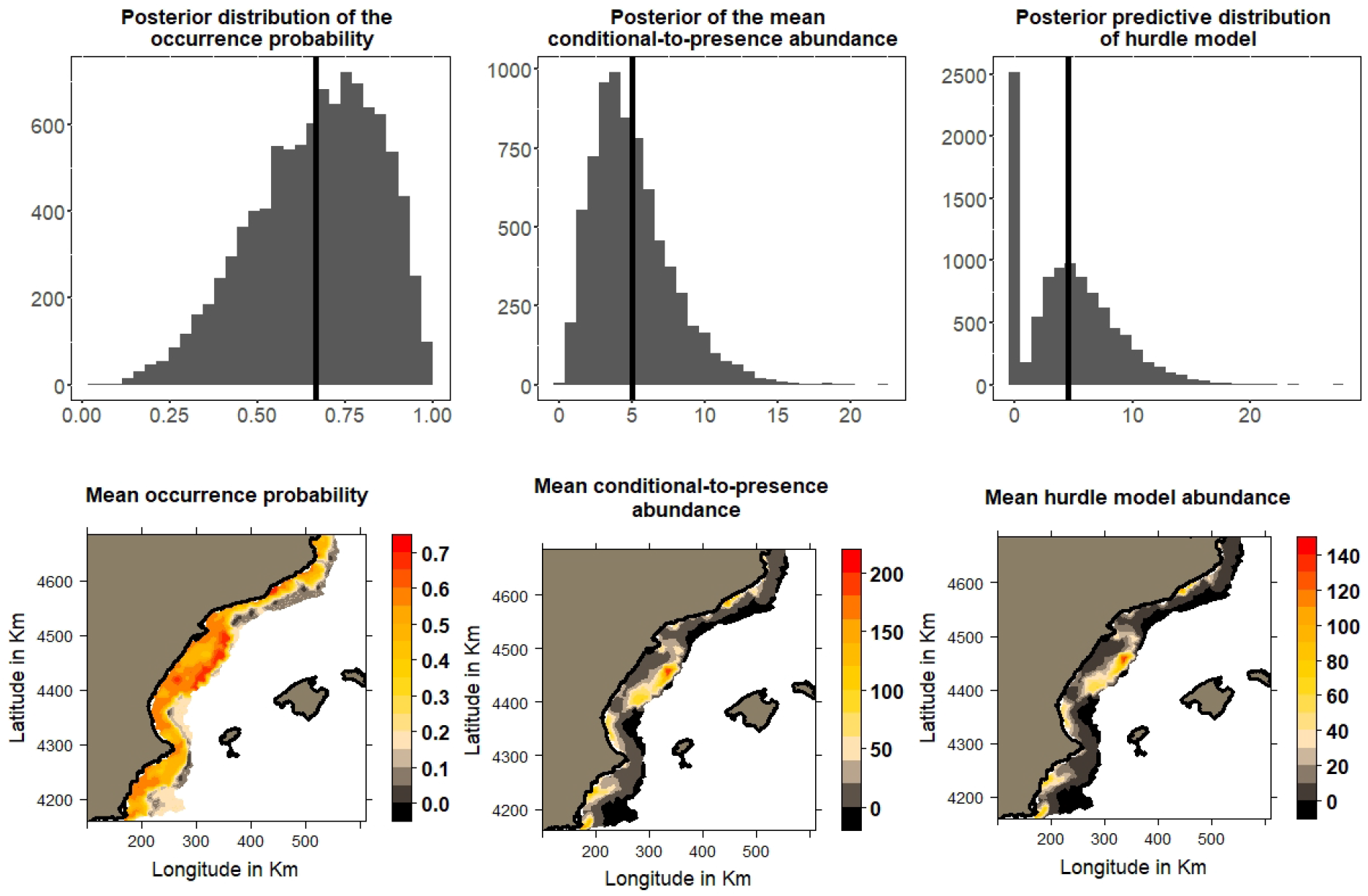
Calculating the predictive posterior distribution of hurdle models using *Mullus barbatus* results for the year 2000. From left to right, top panels represent the mean occurrence probability posterior distribution, the conditional-to-presence mean abundance posterior distribution, and the predictive delta abundance posterior distribution at a particular location. Solid vertical lines represent the mean of the distribution. Bottom panels show mean probability, conditional-to-presence mean abundance, and mean hurdle model abundance maps.

### 2.3 Fisheries management priority area identification

Spatial prioritisation algorithms use several species distribution maps to optimise priority areas based on user defined conservation objectives, constraints, and penalisations. Conservation objectives set the expected ecological targets to meet. Constraints establish a set of prerequisites to the solutions to ensure that solutions exhibit specific properties (e.g. select specific planning units for protection), and penalisations to penalise solutions according to specific metric (e.g. connectivity).

This study identified demersal fisheries management priority areas minimising the size of the area required (using area as a proxy of cost) to protect a minimum set objective, similar to the Marxan’s decision support tool (Ball et al. 2009). The prioritisations were solved to within 1% of optimality using Gurobi (version 8.1.0) (Bixby 2007) and the prioritizr R package (Hanson et al. 2017). We were particularly interested in protecting nursery grounds, so we set our protection targets to 20% of recruits and 10% of non-recruits. We hypothesized that recruits and non-recruits priority areas could be significantly different, thus we fitted three different scenarios based on these targets: one that met both objectives, one that met recruits protection targets alone and another that only met non-recruits protection targets.

In fisheries, fishing effort may be regarded as a proxy to economic impact, and therefore it is common to include Vessel Monitoring System (VMS) or Automatic Identification System (AIS) derived data as a penalisation in the spatial prioritisation algorithm (Afán et al. 2018, Giménez et al. 2020). However, the current overexploitation of fishery resources is driven by an excess of fishery effort (Brochier et al. 2018) and therefore we decided to identify the most productive fishery areas for protection, regardless the impact in the fishery.

#### 2.3.1 Persistence of priority areas

Marine ecosystems and fish assemblages change in space and time (Halpern et al. 2015, Gordon et al. 2018, Kroeker et al. 2020), which has generated arguments in favor of dynamic MPA designs (Grafton & Kompas 2005, Hughes et al. 2007, Hobday & Hartmann 2006). As a result, we assessed the spatiotemporal dynamics of priority areas. To do so, we compared two spatiotemporal optimisations that provided valuable information about the level of spatial persistence of priority areas (See Figure 3). One optimisation included all available maps together (i.e. every time event in the series) to optimise an overall priority area. The overall priority area solution can be regarded as a temporally averaged solution. The other optimisation fitted different priority areas to every time event in the series, providing a temporal series of priority area maps. These maps were then summarised into a frequency map (i.e. a map that overlaps the number of times an area has been selected as a priority area over the time series). The results from these two optimisations were then used to assess the suitability of fixed, progressive or other types of dynamic MPA designs in the study area.

**Figure 3:**
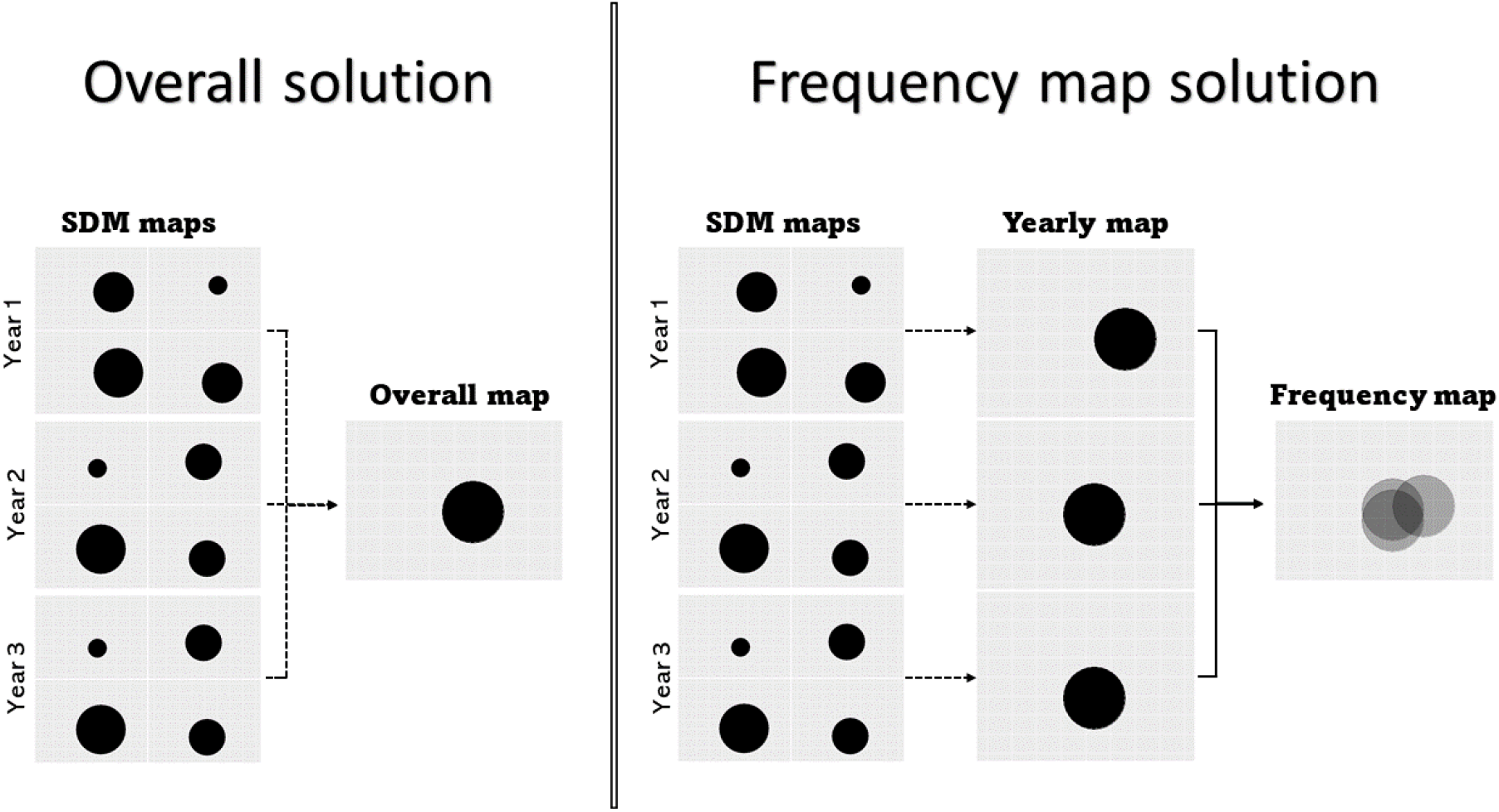
Difference between the overall map (left) and frequency map (right). These results are then used to assess the spatiotemporal dynamics of priority areas. Dashed lines represent the use of a spatial prioritisation algorithm.

##### Persistent priority areas

persistence was assessed by comparing the selected areas in the overall priority area solution and the temporal frequency map solution. High similarity between them implies that yearly priority areas do not differ much from the overally selected priority area, suggesting that priority areas are persistent and therefore a fixed MPA design would be effective. In contrast, divergence in priority areas between the two solutions may suggest a dynamic scenario that requires further considerations and more flexible designs.

##### Progressive priority areas

difference between the overall priority area solution and the temporal frequency map solution suggest some sort of spatiotemporal variability in priority areas. Progressive priority area drifts were assessed using the Cohen’s kappa coefficient, that measure inter-rater reliability for qualitative data (Landis & Koch 1977, McHugh 2012), and help us asses the similarity between different priority area maps (Ban et al. 2009). By doing a pairwise comparison across every map in the time series, we produced a kappa matrix that summarises all the pairwise coefficients and follow the categorisation proposed by Landis & Koch (1977): 0, “No agreement”; 0-0.2, “Slight agreement”; 0.2-0.4, “Fair agreement”; 0.4-0.6, “Moderate agreement”; 0.6-0.8, “Substantial agreement”; and 0.8-1.0, “Almost perfect agreement.” We particularly looked at the diagonal of the kappa matrix because it indicates the similarity between successive yearly priority areas. Therefore, a kappa matrix with consistently high diagonal values manifests a progressive evolution in the distribution of priority areas, and therefore, new MPA designs could be informed progressively based on the last fishery-independent survey data.

##### Other priority area dynamics

inconsistent Cohen’s kappa matrix diagonal values imply more flexible priority area dynamics. Under such a situation, it is useful to identify patterns that help us further understand the spatiotemporal process under study. Whilst humans are able to extract patterns from maps, quantifying similarities and dissimilarities amongst them can be quite challenging, especially when working with several maps. In this regard, we used multivariate methods to help us identify recurrent spatial patterns from the series of maps. Specificaly, we used ordination plots and cluster dendrograms to evaluate similarities between spatial prioritisation solutions (Linke et al. 2011). Non-metric multidimensional scaling procedure (NMDS) allowed us to visualise similarities among different solutions in several dimensions with the advantage that the relative differences between solutions were conserved, so it reflected true dissimilarities. In contrast, clustering methods quantify distance between solutions and allowed us to organise them into a dendrogram to help us choose the number of groups.

## 3 Results

SDM results are summarised in Table 2 and reveal rather persistent distributions for *M. barbatus, M. surmuletus, L. budegassa* recruits and *M. merluccius* recruits and non-recruits, suggesting that the spatial distribution of these species and life stages did not vary much from year to year. *I. coindetii* and *L. piscatorius* recruits and non-recruits showed smooth spatial distribution changes. *L. budegassa* non-recruit distribution changed very abruptly from year to year, falling in the opportunistic distribution category. Figure 4 visualises the average spatial distribution of each species and life stage from 2000 to 2016 (Visit tinyurl.com/42hy9e8m for the full time series of maps). Figure 5 and Figure 6 present the fitted bathymetric distribution and overall population size trends for each species and life stage, respectively.

**Table 2:**
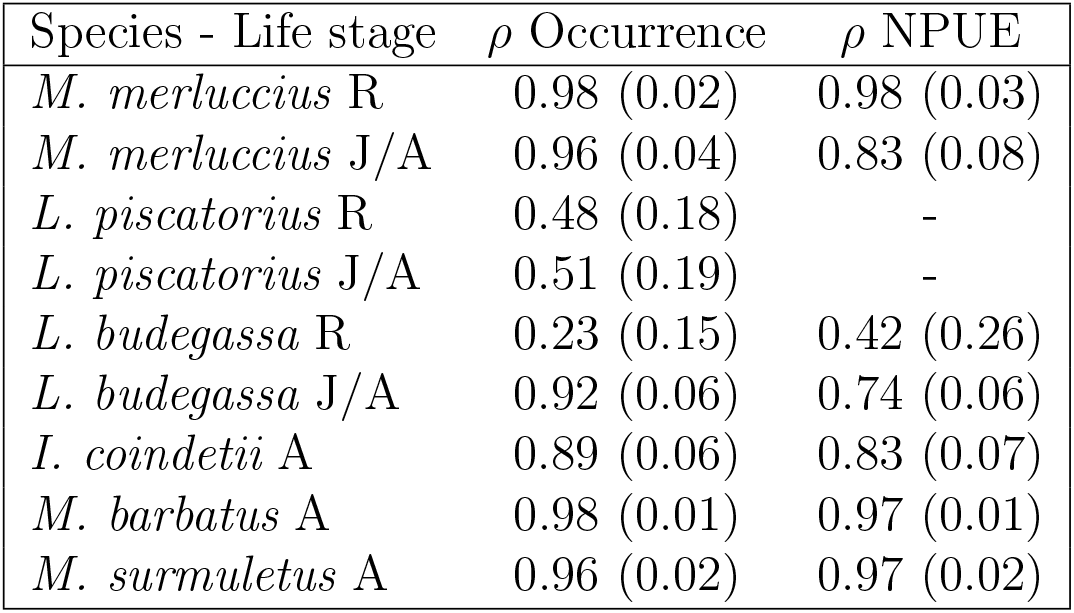
Summary of fitted species-specific spatiotemporal pattern. R and J/A stand for recruits and non-recruits respectively. *ρ* is the temporal autocorrelation parameter of the spatiotemporal structure and the value within the parenthesis is the associated standard deviation.

| Species - Life stage | $\rho$ Occurrence | $\rho$ NPUE |
| --- | --- | --- |
| *M. merluccius* R | 0.98 (0.02) | 0.98 (0.03) |
| *M. merluccius* J/A | 0.96 (0.04) | 0.83 (0.08) |
| *L. piscatorius* R | 0.48 (0.18) | - |
| *L. piscatorius* J/A | 0.51 (0.19) | - |
| *L. budegassa* R | 0.23 (0.15) | 0.42 (0.26) |
| *L. budegassa* J/A | 0.92 (0.06) | 0.74 (0.06) |
| *I. coindetii* A | 0.89 (0.06) | 0.83 (0.07) |
| *M. barbatus* A | 0.98 (0.01) | 0.97 (0.01) |
| *M. surmuletus* A | 0.96 (0.02) | 0.97 (0.02) |

**Figure 4:**
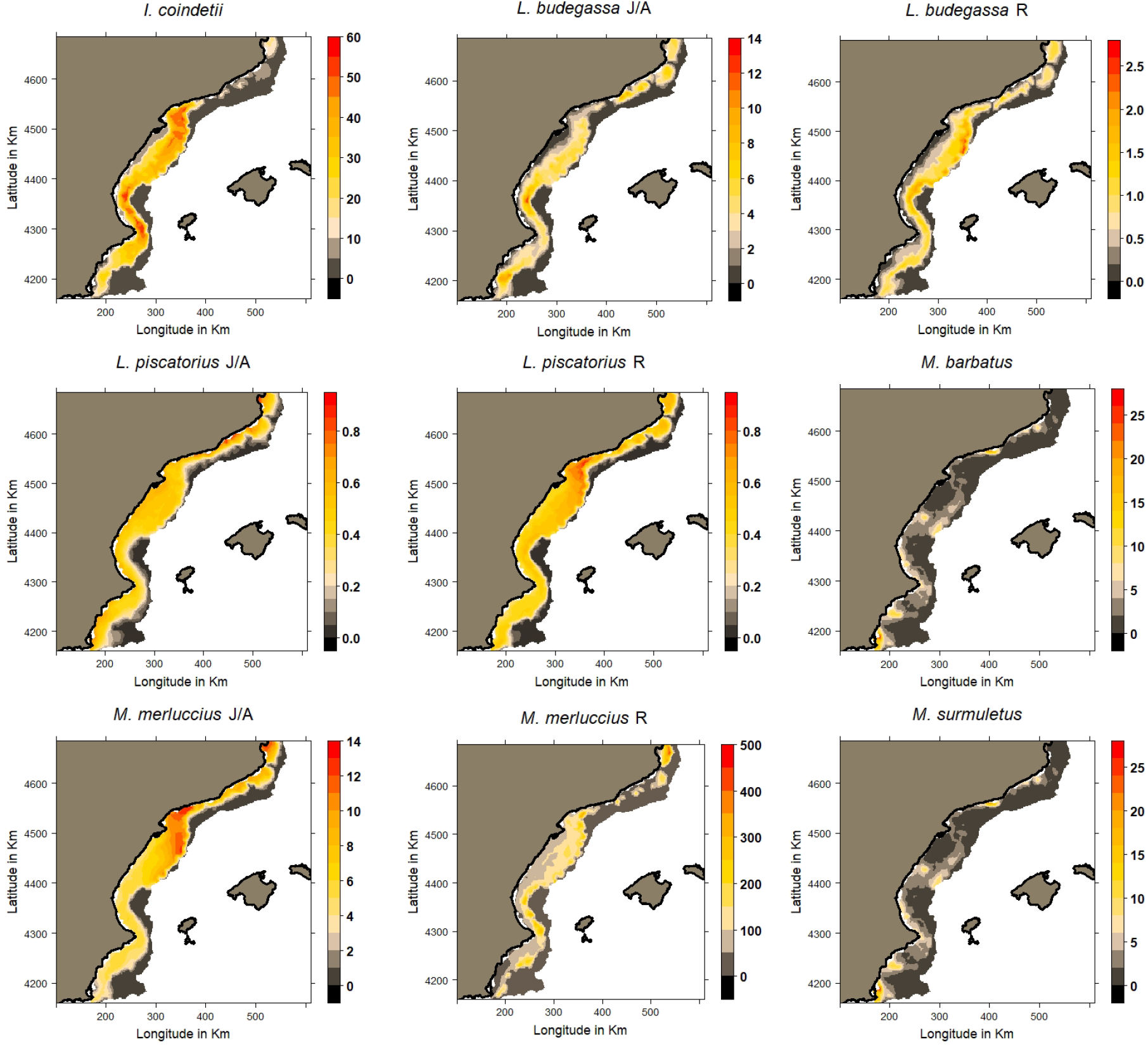
Averaged spatial distributions of the different species and life stage abundances between 2000 and 2016. Visit tinyurl.com/42hy9e8m for yearly maps. R and J/A stand for recruits and non-recruits respectively.

**Figure 5:**
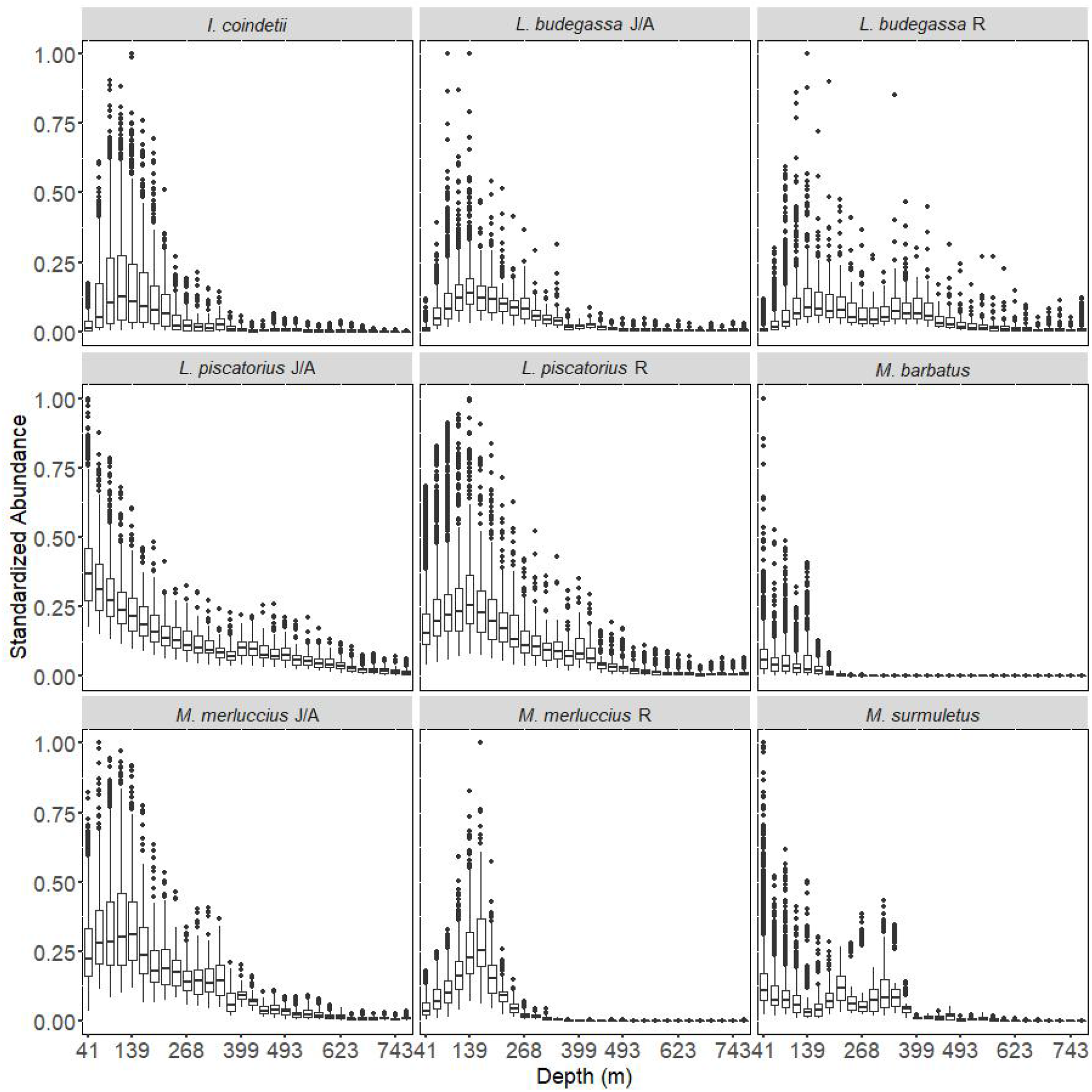
Fitted non-linear bathymetric effects for each of demersal species considered. Y-axis values were standardized to be between 0 and 1 for the sake of comparability. Each boxplot corresponds to an approximately 20 m depth interval. Each box represents the interquartile range of the mean fitted values, the central bold line represents the median value, and dots represent fitted values above 1.5 times and below 3 times the interquartile range beyond either end of the box. R and J/A stand for recruits and non-recruits respectively.

**Figure 6:**
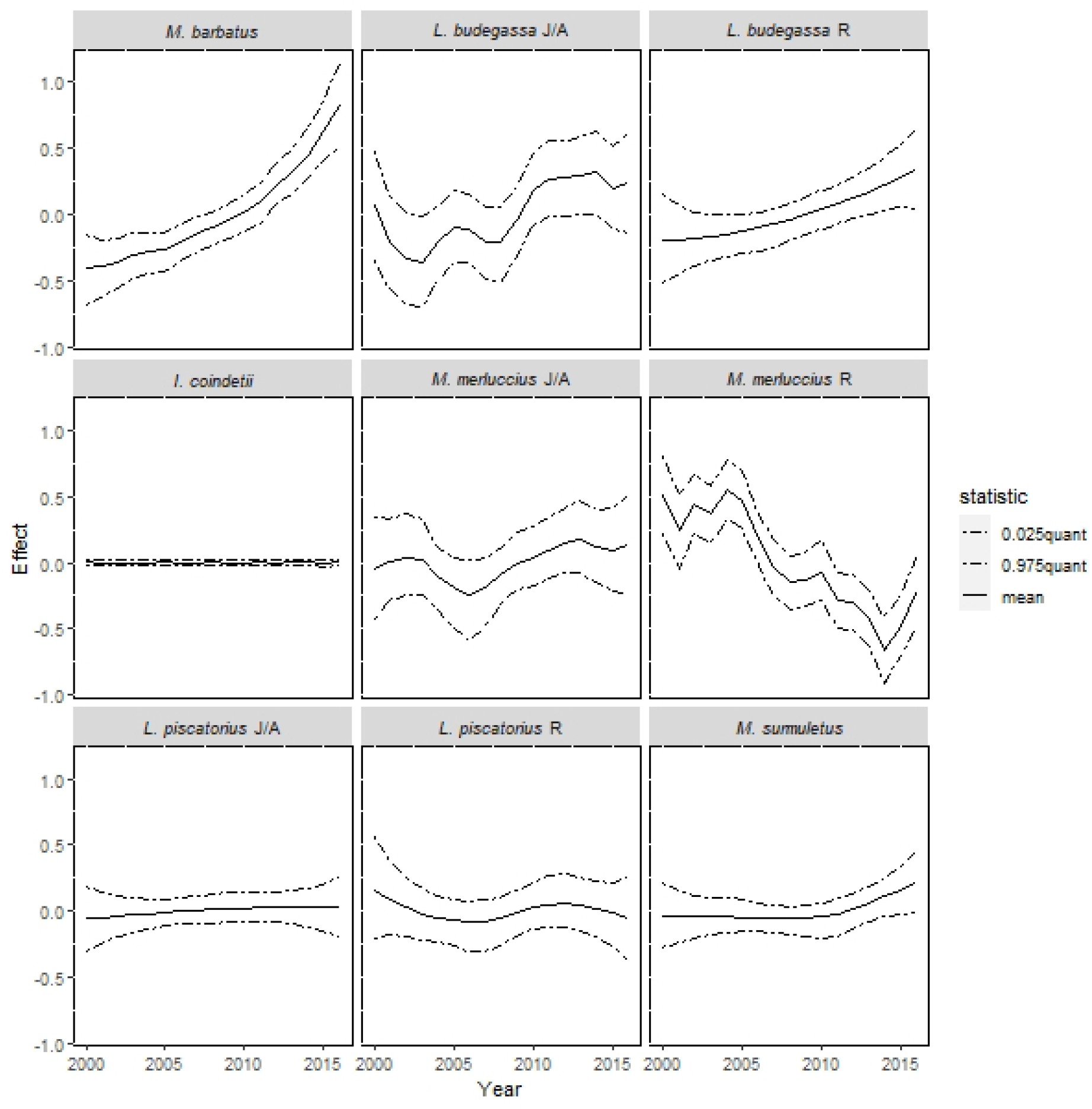
Each species and life stage temporal trends from 2000 to 2016 in the study area. Solid lines represent the mean effect in the linear predictor, and dotted lines represent the 95 credibility interval. R and J/A stand for recruits and non-recruits, respectively.

Figure 7 presents the overall priority areas solution and priority frequency maps for the three different scenarios considered in this study. These results provided two important conclusions for decision making. On the one hand, priority areas that met both recruits and non-recruits protection targets were very similar to the areas that met recruits protection targets alone. In other words, by optimising the protection of 20% of recruits, we also protected 10% of non-recruits in the study area. As a consequence, from this point forward, the study focused on the scenario that combined the 20% recruits and 10% non-recruits protection target altogether. On the other hand, the overall priority area solution and yearly priority area solution were substantially different implying non-persistent priority areas during the study period. Therefore, we deduced that a fixed MPAs design could be ineffective and decided to further explore the spatiotemporal dynamics of priority areas.

**Figure 7:**
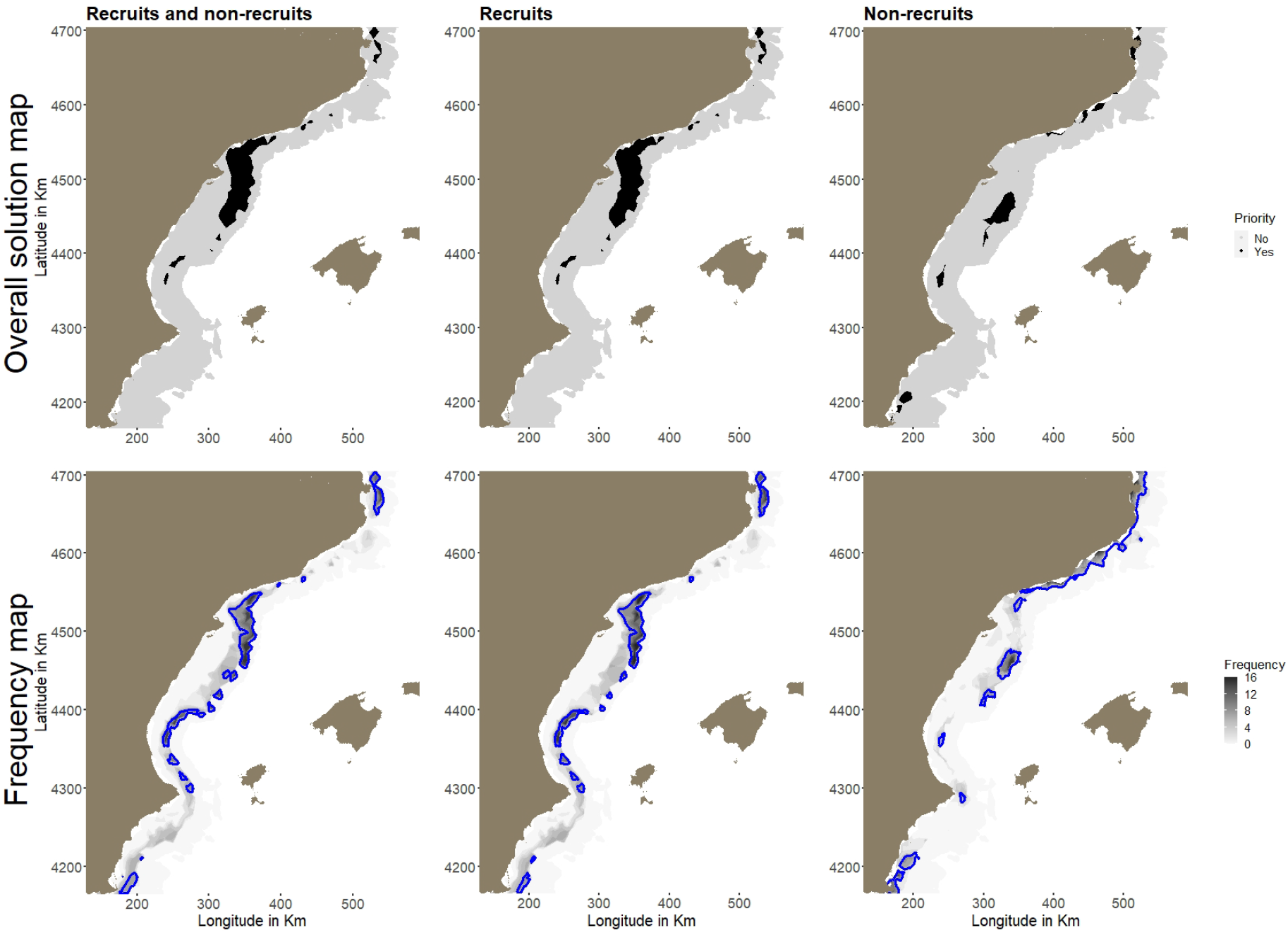
Overall priority areas and priority area frequency maps for the different conservation targets. Blue lines represent the 90^*th*^ percentile contour lines.

Cohen’s kappa matrix diagonal indicated that the similarity between temporally subsequent priority areas was rather inconsistent (Figure 8). Therefore, we deduced that a progressive MPA design based on the previous year’s survey result would not be very effective.

**Figure 8:**
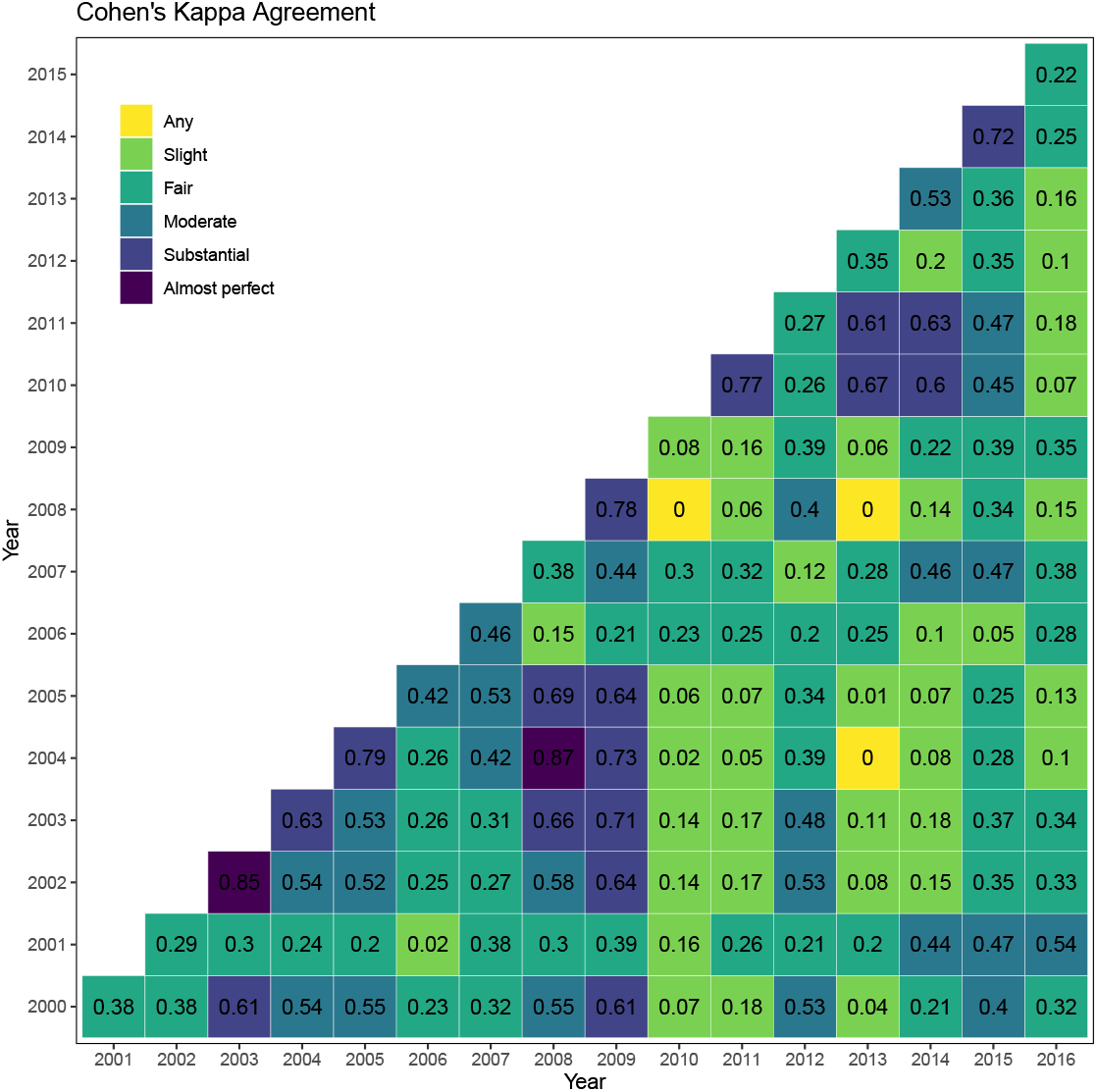
Cohen’s kappa statistic matrix comparing pairwise yearly priority area maps. Of special interest is the diagonal of the kappa matrix, which indicates the similarity between temporally subsequent priority areas. A scenario where kappa matrix diagonal values are consistently high manifests a progressive evolution in the priority areas.

NMDS and hierarchical clustering results suggested the presence of either two or four patterns (Figure 9). After carefully looking at the solutions and getting the opinion of fishery experts’, we decided to select two groups as highlighted by the different colors. Interestingly, there seems to be a clear temporal pattern between these two clusters, one cluster occurring during the first part of the series and the other one in the second part (See right panel in Figure 9). We used these results to create new frequency maps based on these two clusters.

**Figure 9:**
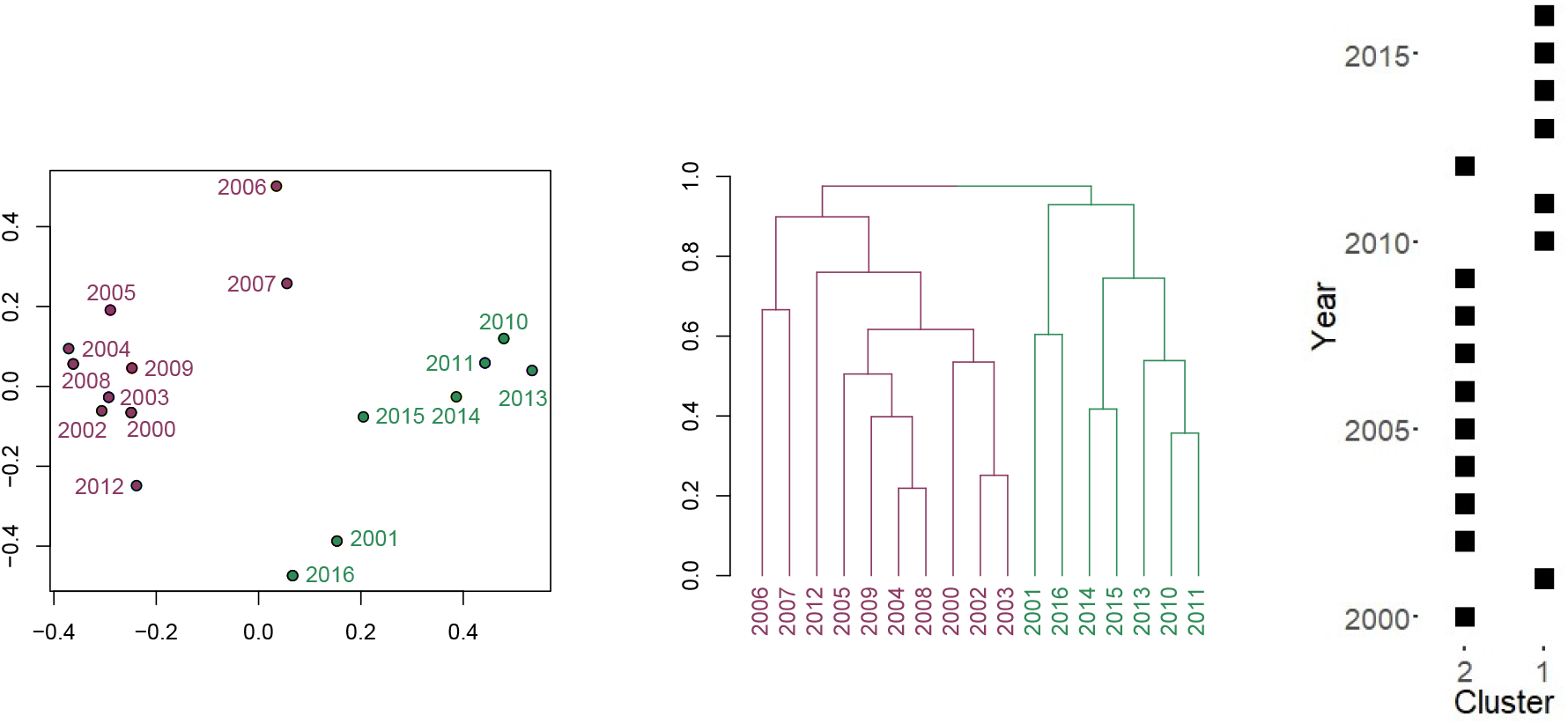
NMDS scatterplot (left panel) and hierarchical clustering dendrogram (centre panel) of yearly priority area results. The right panel shows the time series of the selected clusters.

In the end, we obtained a portfolio of priority area maps (Figure 10) containing: an overall solution map (top-left panel); an overall frequency map (top-right panel); and a set of frequency maps for the selected number of clusters (bottom panels). The different maps in Figure 10 consistently identified a number of priority areas. From north to south: 1) the area around Cap de Creus; 2) the end of the continental shelf around the Ebro delta; and 3) the shelf break in front of Valencia. Lastly, despite the lower consistency of priority areas in the south, the zone in front of Mar Menor seemed to be a relatively important ecological area during the second half of the time series (Cluster 1).

**Figure 10:**
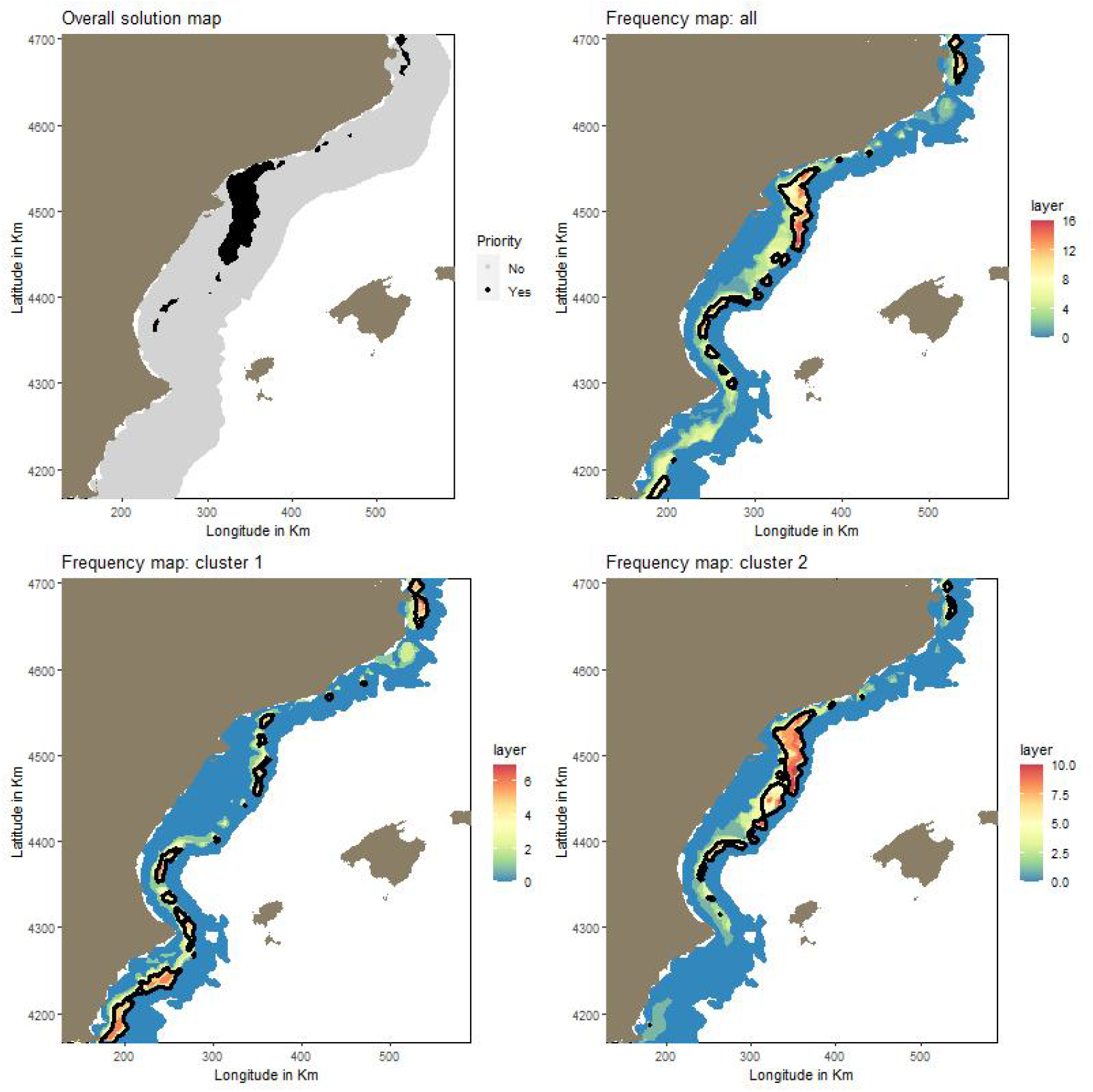
Portfolio of proposed solutions to assess the spatiotemporal distribution of multispecies hotspots as described in Section 2.3.1. The top-left and top-right panels represent the overall solution map and the overall priority area frequency map respectively. The bottom panels show the clustered frequency maps.

## 4 Discussion

This study assessed the spatiotemporal dynamics of fisheries priority areas using: spatiotemporal SDMs; different spatiotemporal prioritisation configurations; and a small set of simple metrics and multivariate methods. The method was applied over a western Mediterranean case study and results identified two temporally structured priority area patterns, one occurring during the first half of the time series and another during the second half. Identifying the drivers behind these patterns was beyond the scope of this study, but could indicate the presence of a large scale driver worth investigating.

Species distribution maps represent the baseline unit for the identification of fisheries priority areas. Therefore, good spatiotemporal SDMs are essential to adequately asssess the spatiotemporal dynamics of fisheries priority areas. There is a wide range of spatiotemporal SDM software available in `R` (R Core Team 2021). In fisheries, `INLA` (Cosandey-Godin et al. 2015, Paradinas et al. 2020), `VAST` (Thorson 2019) and `mgcv` (Schmiing et al. 2013, Parra et al. 2017) may be the most widely used R packages in fisheries. We provide R scripts (`link`) to fit generic spatiotemporal hurdle SDMs using INLA so that other users may apply them in other case studies and areas of interest.

Our spatial prioritisation optimisations did not include fishery effort as a penalization and identified fisheries priority areas based on conservation objectives alone. Far form considering the impact on fishers unimportant, given the overexploitation levels in the Mediterranean (FAO 2000, Sola et al. 2020), our objective was to identify the most important areas for the species under study. In fact, the socioeconomic impact may be assessed afterwards by calculating the amount of fishing activity that would need to be relocated in other areas using Vessel Monitoring System (VMS) or Automatic Identification System (AIS) derived data.

Our case study constitutes a clear example where fishery priority areas change in space and time. Non spatiotemporal SDMs and/or prioritisations would have ignored such variability, potentially reducing the conservation impact. In this regard, long-term surveys are essential to predict the spatiotemporal distribution of species, and assess the spatiotemporal variability of priority areas. Similarly, intra-annual temporal resolution is key to assess year around persistence of priority areas. Unfortunately, fishery-independent surveys are generally programmed once or twice a year, and identified conservation priority areas may not be representative of all seasons. Fishery dependent data could complement survey data, but its spatial coverage is not always scalable to survey data and target species estimates are affected by the preferential sampling bias (Diggle et al. 2010, Pennino et al. 2018). Another relatively cheap option to complement fishery-independent surveys could be to seek the collaboration of the fishery sector to scientifically sample the ocean in different seasons. The Norwegian reference fleet (Nedreaas et al. 2006) constitutes an excellent example of co-operation between fishers and scientists.

The described procedure follows a clear step-by-step approach to assess the spatiotemporal dynamics of conservation priority areas. A good implementation requires: highquality and long-term fishery data; on-site fishery knowledge to select key conservation objectives; expertise to fit robust spatiotemporal SDMs; spatial planning software skills; and basic multivariate analysis understanding. While this method has been developed for fisheries management, it is also applicable to other multispecies systems that evolve with time, no matter marine or terrestrial. Clearly, there is still a considerable challenge ahead to collect quality fishery data to inform the intra-annual dynamics of conservation priority areas. Lastly, we would like to suggest that, given the spatiotemporal dynamics of marine systems and fishery markets, existing MPA designs should go through cyclical and iterative re-assessments that incorporate new information and adapt their objectives and measures according to the evolution of the socio-ecological system.

## 5 Acknowledgements

The authors express their gratitude to all the people that work in the MEDITS surveys and especially to Antonio Esteban for his invaluable collaboration. MEDITs project has been co-funded since 1994 by the EU through the European Maritime and Fisheries Fund (EMFF) within the National Program of collection, management and use of data in the fisheries sector and support for scientific advice regarding the Common Fisheries Policy. I.P. would like to thank the Fundación Biodiversidad (Spanish Ministry of Agriculture and the Environment) for the funding provided and the invaluable help provided to access the required data. D.C. and A.L.Q. also thank the Spanish Ministerio de Ciencia e Innovación — Agencia Estatal de Investigación for grant PID2019-106341GB-I00 (jointly financed by the European Regional Development Fund, FEDER). M.G.P. also thanks the IMPRESS (RTI2018-099868-B-I00) project, ERDF, Ministry of Science, Innovation and Universities - State Research Agency. J.G. was supported by the Spanish National Program Juan de la Cierva-Formación (FJC2019-040016-I). This work acknowledges the ‘Severo Ochoa Centre of Excellence’ accreditation (CEX2019-000928-S) to the Institute of Marine Science (ICM-CSIC).

## Notes

### Competing Interest Statement

The authors have declared no competing interest.

### Summary of Updates

This new version is a summarised version of the original manuscript. In light of criticism in Fish and Fisheries due to its rather report like structure, the new manuscript is more focused on the unexpected results, and less in the method. The method is still carefully described but not as extensively as before.

## References

Afán, I., Giménez, J., Forero, M. & Ramírez, F. (2018), ‘An adaptive method for identifying marine areas of high conservation priority’, Conservation Biology 32(6), 1436–1447.

Baillie, J. & Zhang, Y.-P. (2018), ‘Space for nature’.

Ball, I. & Possingham, H. (2000), ‘Marxan (v1. 8.2)’, Marine Reserve Design Using Spatially Explicit Annealing, a Manual.

Ball, I. R., Possingham, H. P. & Watts, M. (2009), ‘Marxan and relatives: software for spatial conservation prioritisation’, Spatial conservation prioritisation: Quantitative methods and computational tools pp. 185–195.

Ban, N. C., Picard, C. R. & Vincent, A. C. (2009), ‘Comparing and integrating community-based and science-based approaches to prioritizing marine areas for protection’, Conservation Biology 23(4), 899–910.

Bertrand, J. A., de Sola, L. G., Papaconstantinou, C., Relini, G. & Souplet, A. (2002), ‘The general specifications of the medits surveys’, Scientia marina 66(S2), 9–17.

Bixby, B. (2007), ‘The gurobi optimizer’, Transp. Re-search Part B 41(2), 159–178.

Brochier, T., Auger, P., Thiao, D., Bah, A., Ly, S., Nguyen-Huu, T. & Brehmer, P. (2018), ‘Can overexploited fisheries recover by self-organization? reallocation of fishing effort as an emergent form of governance’, Marine Policy 95, 46–56.

Cosandey-Godin, A., Krainski, E. T., Worm, B. & Flemming, J. M. (2015), ‘Applying bayesian spatiotemporal models to fisheries bycatch in the canadian arctic’, Canadian Journal of Fisheries and Aquatic Sciences 72(2), 186–197.

Di Franco, A., Thiriet, P., Di Carlo, G., Dimitriadis, C., Francour, P., Gutiérrez, N. L., de Grissac, A. J., Koutsoubas, D., Milazzo, M., del Mar Otero, M. et al. (2016), ‘Five key attributes can increase marine protected areas performance for small-scale fisheries management’, Scientific reports 6(1), 1–9.

Di Lorenzo, M., Claudet, J. & Guidetti, P. (2016), ‘Spillover from marine protected areas to adjacent fisheries has an ecological and a fishery component’, Journal for Nature Conservation 32, 62–66.

Diggle, P. J., Menezes, R. & Su, T. (2010), ‘Geostatistical inference under preferential sampling’, Journal of the Royal Statistical Society: Series C (Applied Statistics) 59(2), 191–232.

Dinerstein, E., Vynne, C., Sala, E., Joshi, A. R., Fernando, S., Lovejoy, T. E., Mayorga, J., Olson, D., Asner, G., Baillie, J. et al. (2019), ‘A global deal for nature: guiding principles, milestones, and targets’, Science advances 5(4), 2869.

Fahrmeir, L. & Lang, S. (2001), ‘Bayesian inference for generalized additive mixed models based on markov random field priors’, Journal of the Royal Statistical Society: Series C (Applied Statistics) 50(2), 201–220.

FAO (2000), The State of World Fisheries and Aquaculture, Vol. 3, Food & Agriculture Org.

Fraschetti, S., Pipitone, C., Mazaris, A. D., Rilov, G., Badalamenti, F., Bevilacqua, S., Claudet, J., Carić, H., Dahl, K., D’Anna, G. et al. (2018), ‘Light and shade in marine conservation across european and contiguous seas’, Frontiers in marine science 5, 420.

Fuglstad, G. A., Simpson, D., Lindgren, F. & Rue, H. (2019), ‘Constructing priors that penalize the complexity of Gaussian random fields’, Journal of the American Statistical Association 114(525), 445–452.

Game, E. T., Bode, M., McDonald-Madden, E., Grantham, H. S. & Possingham, H. P. (2009), ‘Dynamic marine protected areas can improve the resilience of coral reef systems’, Ecology Letters 12(12), 1336–1346.

Giakoumi, S., Scianna, C., Plass-Johnson, J., Micheli, F., Grorud-Colvert, K., Thiriet, P., Claudet, J., Di Carlo, G., Di Franco, A., Gaines, S. D. et al. (2017), ‘Ecological effects of full and partial protection in the crowded mediterranean sea: a regional meta-analysis’, Scientific reports 7(1), 1–12.

Giménez, J., Cardador, L., Mazor, T., Kark, S., Bellido, J. M., Coll, M. & Navarro, J. (2020), ‘Marine protected areas for demersal elasmobranchs in highly exploited mediterranean ecosystems’, Marine Environmental Research p. 105033.

Gordon, T., Harding, H., Clever, F., Davidson, I., Davison, W., Montgomery, D., Weatherhead, R., Windsor, F., Armstrong, J., Bardonnet, A. et al. (2018), ‘Fishes in a changing world: learning from the past to promote sustainability of fish populations’, Journal of fish biology 92(3), 804–827.

Grafton, R. Q. & Kompas, T. (2005), ‘Uncertainty and the active adaptive management of marine reserves’, Marine Policy 29(5), 471–479.

Halpern, B. S., Frazier, M., Potapenko, J., Casey, K. S., Koenig, K., Longo, C., Lowndes, J. S., Rockwood, R. C., Selig, E. R., Selkoe, K. A. et al. (2015), ‘Spatial and temporal changes in cumulative human impacts on the world’s ocean’, Nature communications 6(1), 1–7.

Hanson, J. O., Fuller, R. A. & Rhodes, J. R. (2019), ‘Conventional methods for enhancing connectivity in conservation planning do not always maintain gene flow’, Journal of Applied Ecology 56(4), 913–922.

Hanson, J., Schuster, R., Morrell, N., Strimas-Mackey, M., Watts, M., Arcese, P. & Possingham, H. (2017), ‘prioritizr: systematic conservation prioritization in r’, R package.(version 1.0. 1.0 ed).

Hilborn, R., Stokes, K., Maguire, J.-J., Smith, T., Botsford, L. W., Mangel, M., Orensanz, J., Parma, A., Rice, J., Bell, J. et al. (2004), ‘When can marine reserves improve fisheries management?’, Ocean & Coastal Management 47(3-4), 197–205.

Hobday, A. & Hartmann, K. (2006), ‘Near real-time spatial management based on habitat predictions for a longline bycatch species’, Fisheries Management and Ecology 13(6), 365–380.

Hughes, T. P., Gunderson, L. H., Folke, C., Baird, A. H., Bellwood, D., Berkes, F., Crona, B., Helfgott, A., Leslie, H., Norberg, J. et al. (2007), ‘Adaptive management of the great barrier reef and the grand canyon world heritage areas’, AMBIO: A Journal of the Human Environment 36(7), 586–592.

Izquierdo, F., Paradinas, I., Cerviño, S., Conesa, D., Alonso-Fernández, A., Velasco, F., Preciado, I., Punzón, A., Saborido-Rey, F. & Pennino, M. G. (2021), ‘Spatiotemporal assessment of the european hake (merluccius merluccius) recruits in the northern iberian peninsula’, Frontiers in Marine Science 8, 1.

Kaiser, M. J., Blyth-Skyrme, R. E., Hart, P. J., Edwards-Jones, G. & Palmer, D. (2007), ‘Evidence for greater reproductive output per unit area in areas protected from fishing’, Canadian Journal of Fisheries and Aquatic Sciences 64(9), 1284–1289.

Kerwath, S. E., Winker, H., Götz, A. & Attwood, C. G. (2013), ‘Marine protected area improves yield without disadvantaging fishers’, Nature communications 4(1), 1–6.

Kroeker, K. J., Bell, L. E., Donham, E. M., Hoshijima, U., Lummis, S., Toy, J. A. & Willis-Norton, E. (2020), ‘Ecological change in dynamic environments: Accounting for temporal environmental variability in studies of ocean change biology’, Global Change Biology 26(1), 54–67.

Landis, J. R. & Koch, G. G. (1977), ‘The measurement of observer agreement for categorical data’, biometrics pp. 159–174.

Lang, S. & Brezger, A. (2004), ‘Bayesian p-splines’, Journal of computational and graphical statistics 13(1), 183–212.

Lecomte, J.-B., Benôit, H. P., Ancelet, S., Etienne, M.-P., Bel, L. & Parent, E. (2013), ‘Compound poisson-gamma vs. delta-gamma to handle zero-inflated continuous data under a variable sampling volume’, Methods in Ecology and Evolution 4(12), 1159–1166.

Leenhardt, P., Low, N., Pascal, N., Micheli, F. & Claudet, J. (2015), The role of marine protected areas in providing ecosystem services, in ‘Aquatic functional biodiversity’, Elsevier, pp. 211–239.

Lindgren, F., Rue, H. & Lindström, J. (2011), ‘An explicit link between gaussian fields 670 and gaussian markov random fields: the spde approach (with discussion)’, JR 671 Stat Soc, Series B 73, 423–498.

Linke, S., Watts, M., Stewart, R. & Possingham, H. P. (2011), ‘Using multivariate analysis to deliver conservation planning products that align with practitioner needs’, Ecography 34(2), 203–207.

Lizaso, J. L. S., Sola, I., Guijarro-García, E., Bellido, J. M. & Franquesa, R. (2020), ‘A new management framework for western mediterranean demersal fisheries’, Marine Policy 112, 103772.

Martín, P., Maynou, F., Garriga-Panisello, M., Ramírez, J. & Recasens, L. (2019), ‘Fishing effort alternatives for the management of demersal fisheries in the western mediterranean’, Scientia Marina 83(4), 293–304.

Martin, T. G., Wintle, B. A., Rhodes, J. R., Kuhnert, P. M., Field, S. A., Low-Choy, S. J., Tyre, A. J. & Possingham, H. P. (2005), ‘Zero tolerance ecology: improving ecological inference by modelling the source of zero observations’, Ecology letters 8(11), 1235–1246.

Martínez-Minaya, J., Cameletti, M., Conesa, D. & Pennino, M. G. (2018), ‘Species distribution modeling: a statistical review with focus in spatio-temporal issues’, Stochastic environmental research and risk assessment 32(11), 3227–3244.

Martins, T. G., Simpson, D., Lindgren, F. & Rue, H. (2013), ‘Bayesian computing with inla: new features’, Computational Statistics & Data Analysis 67, 68–83.

Maunder, M. N. & Punt, A. E. (2004), ‘Standardizing catch and effort data: a review of recent approaches’, Fisheries research 70(2-3), 141–159.

Maynou, F. (2014), ‘Coviability analysis of western mediterranean fisheries under msy scenarios for 2020’, ICES Journal of Marine Science 71(7), 1563–1571.

McHugh, M. L. (2012), ‘Interrater reliability: the kappa statistic’, Biochemia medica: Biochemia medica 22(3), 276–282.

Moilanen, A., Kujala, H. & Leathwick, J. (2009), ‘The zonation framework and software for conservation prioritization’, Spatial conservation prioritization 135, 196–210.

Nedreaas, K. H., Borge, A., Godøy, H. & Aanes, S. (2006), The norwegian reference fleet: co-operation between fisherman and scientists for multiple objectives, ICES.

O’Leary, B. C., Winther-Janson, M., Bainbridge, J. M., Aitken, J., Hawkins, J. P. & Roberts, C. M. (2016), ‘Effective coverage targets for ocean protection’, Conservation Letters 9(6), 398–404.

Ovando, D., Dougherty, D. & Wilson, J. R. (2016), ‘Market and design solutions to the short-term economic impacts of marine reserves’, Fish and Fisheries 17(4), 939–954.

Paradinas, I., Conesa, D., López-Quílez, A. & Bellido, J. M. (2017), ‘Spatio-temporal model structures with shared components for semi-continuous species distribution modelling’, Spatial Statistics 22, 434–450.

Paradinas, I., Conesa, D., López-Quílez, A., Esteban, A., López, L. M. M., Bellido, J. M. & Pennino, M. G. (2020), ‘Assessing the spatiotemporal persistence of fish distributions: a case study on two red mullet species (mullus surmuletus and m. barbatus) in the western mediterranean’, Marine Ecology Progress Series 644, 173–185.

Parra, H. E., Pham, C. K., Menezes, G. M., Rosa, A., Tempera, F. & Morato, T. (2017), ‘Predictive modeling of deep-sea fish distribution in the azores’, Deep Sea Research Part II: Topical Studies in Oceanography 145, 49–60.

Pennino, M. G., Paradinas, I., Illian, J. B., Muñoz, F., Bellido, J. M., López-Quílez, A. & Conesa, D. (2018), ‘Accounting for preferential sampling in species distribution models’, Ecology and Evolution.

Petza, D., Anastopoulos, P., Coll, M., Garcia, S., Kaiser, M., Kalogirou, S., Lourdi, I., Rice, J., Sciberras, M. & Katsanevakis, S. (2021), ‘The contribution of area-based fisheries management measures to fisheries sustainability and marine conservation: a global scoping review protocol’, Research Ideas and Outcomes 7, e70486.

Petza, D., Maina, I., Koukourouvli, N., Dimarchopoulou, D., Akrivos, D., Kavadas, S., Tsikliras, A., Karachle, P. & Katsanevakis, S. (2017), ‘Where not to fish–reviewing and mapping fisheries restricted areas in the aegean sea’, Mediterranean Marine Science 18(2), 310–323.

R Core Team (2021), R: A Language and Environment for Statistical Computing, R Foundation for Statistical Computing, Vienna, Austria. URL: https://www.R-project.org/

Raicevich, S., Alegret, J.-L., Frangoudes, K., Giovanardi, O. & Fortibuoni, T. (2018), ‘Community-based management of the mediterranean coastal fisheries: Historical reminiscence or the root for new fisheries governance?’, Regional Studies in Marine Science 21, 86–93.

Rodríguez, J. P., Brotons, L., Bustamante, J. & Seoane, J. (2007), ‘The application of predictive modelling of species distribution to biodiversity conservation’, Diversity and Distributions pp. 243–251.

Schmiing, M., Afonso, P., Tempera, F. & Santos, R. S. (2013), ‘Predictive habitat modelling of reef fishes with contrasting trophic ecologies’, Marine Ecology Progress Series 474, 201–216.

Seo, C., Thorne, J. H., Hannah, L. & Thuiller, W. (2009), ‘Scale effects in species distribution models: implications for conservation planning under climate change’, Biology letters 5(1), 39–43.

Simpson, D., Rue, H., Riebler, A., Martins, T. G. & Sørbye, S. H. (2017), ‘Penalising model component complexity: A principled, practical approach to constructing priors’, Statistical science pp. 1–28.

Sola, I., Maynou, F. & Sánchez Lizaso, J.L. (2020), ‘Bioeconomic analysis of the eu multiannual management plan for demersal fisheries in the western mediterranean. spanish fisheries as a case study’, Frontiers in Marine Science.

Stefánsson, G. (1996), ‘Analysis of groundfish survey abundance data: combining the glm and delta approaches’, ICES journal of Marine Science 53(3), 577–588.

Thorson, J. T. (2019), ‘Guidance for decisions using the vector autoregressive spatiotemporal (vast) package in stock, ecosystem, habitat and climate assessments’, Fisheries Research 210, 143–161.

Tittensor, D. P., Walpole, M., Hill, S. L., Boyce, D. G., Britten, G. L., Burgess, N. D., Butchart, S. H., Leadley, P. W., Regan, E. C., Alkemade, R. et al. (2014), ‘A mid-term analysis of progress toward international biodiversity targets’, Science 346(6206), 241–244.

Zuur, A. F., Ieno, E. N., Walker, N. J., Saveliev, A. A. & Smith, G. M. (2009), Zero-truncated and zero-inflated models for count data, in ‘Mixed effects models and extensions in ecology with R’, Springer, pp. 261–293.

